# Genomic regions associated with *Mycoplasma ovipneumoniae* presence in nasal secretions of domestic sheep

**DOI:** 10.1101/2021.02.04.429710

**Authors:** Michelle R. Mousel, Stephen N. White, Maria K. Herndon, David R. Herndon, J. Bret Taylor, Gabrielle M. Becker, Brenda M. Murdoch

## Abstract

*Mycoplasma ovipneumoniae* contributes to polymicrobial pneumonia in domestic sheep. Elucidation of host genetic components to *M. ovipneumoniae* nasal detection would have the potential to reduce the incidence of pneumonia. Nasal mucosal secretions were collected from 647 sheep from a large US sheep flock. Ewes of three breeds (Polypay n=222, Rambouillet n=321, and Suffolk n=104) ranging in age from one to seven years, were sampled at three different times of the production cycle (February, April, and September/October) over four years (2015 to 2018). The presence and quantity of *M. ovipneumoniae* was determined using a species-specific qPCR with 3 to 10 sampling times per sheep. Breed (P<0.001), age (P<0.024), sampling time (P<0.001), and year (P<0.001) of collection affected log_10_ transformed *M. ovipneumoniae* DNA copy number, where Rambouillet had the lowest (P<0.0001) compared with both Polypay and Suffolk demonstrating a possible genetic component to detection. Samples from yearlings, April, and 2018 had the highest (P<0.046) detected DNA copy number mean. Sheep genomic DNA was genotyped with the Illumina OvineHD BeadChip. Principal component analysis identified most of the variation in the dataset was associated with breed. Therefore, genome wide association analysis was conducted with a mixed model (EMMAX), with principle components 1 to 6 as fixed and a kinship matrix as random effects. Genome-wide significant (P<9×10^−8^) SNPs were identified on chromosomes 6 and 7 in the all-breed analysis. Individual breed analysis had genome-wide significant (P<9×10^−8^) SNPs on chromosomes 3, 4, 7, 9, 10, 15, 17, and 22. Annotated genes near these SNPs are part of immune (*ANAPC7, CUL5, TMEM229B, PTPN13*), gene translation (*PIWIL4*), and chromatin organization (*KDM2B*) pathways. Immune genes are expected to have increased expression when leukocytes encounter *M. ovipneumoniae* which would lead to chromatin reorganization. Work is underway to narrow the range of these associated regions to identify the underlying causal mutations.

## Introduction

*Mycoplasma ovipneumoniae* is considered a commensal bacterium [1] found in domesticated sheep and goats, but it can at times play a role in the aetiology of chronic, non-progressive pneumonia [2,3]. The bacteria adhere to and infect airway epithelial cells and can cause oxidative damage and apoptosis through an *extracellular-signal-regulated kinase* signaling-mediated mitochondrial pathway [4]. In addition, *M. ovipneumoniae* can moderate the host immune system by eliciting alterations in macrophage activity [5], suppress lymphocytes [6], and induce production of autoantibodies to host ciliary antigen [7].

At slaughter, 83 to 90% of lambs’ lungs with chronic bronchopneumonia were positive for *M. ovipneumoniae* [8,9]. In the U.S., 88.5% of flocks had individual sheep with *M. ovipneumoniae* detected in their nasal secretions (APHIS Info Sheet #708.0615). Using National Animal Health Monitoring System data, comparison of positive with negative *M. ovipneumoniae* flocks revealed a slight decrease in productivity but known effects that could contribute to this small difference (∼4% [10]), such as litter size and ewe age, were not considered in the comparison. Seroprevalence of *M. ovipneumoniae* was found to be highly variable (9.76 to 30.61%) among Kazak, Hu, Merino, and Duolong sheep breeds [11]. This breed variability suggested there may be a genetic component that is impacting bacterial shedding [11,12]. Genome-wide association studies (GWAS) can identify associated genomic regions of interest with underlying causal mutations that affect infectious disease traits.

Technologies have been developed to improve the probability of identifying genomic regions associated with phenotypic traits of interest in sheep. The Ovine SNP50 beadchip [13] and the OvineHD BeadChip (600K) were collaboratively, internationally developed and have been used to identify new markers associated with inherited diseases [14–16], erythrocyte traits [17], parasite infection [18–20], and other infectious disease traits [21–24].

Previous exposure to *M. ovipneumoniae* can be assessed by the presence of antibodies [11], though this does not enable quantification of bacteria equivalents present. There are several molecular amplification assays used to detect this bacterium [25–27]. In order to eliminate false positives and ensure bacterium-specific quantification of *M. ovipneumoniae*, a species-specific quantitative PCR was developed for use with DNA from nasal secretions. Previous work utilizing a modified PCR [28] for screening of domestic animals and wildlife resulted in unintended or non-specific amplification of other bacteria (unpublished data). The assay used in this study permitted a more precise phenotype measurement.

The objective of this study was to quantify *M. ovipneumoniae* in three prevalent US sheep breeds over time and then identify genomic regions associated with *M. ovipneumoniae* detection. These breeds were developed and continue to be selected for high quality wool, fast growing, muscular lambs, or larger litters with good mothering ability to ensure wide applicability of the findings for the sheep industry. This study elucidated *M. ovipneumoniae* detection patterns over time and identified genomic regions associated with detected *M. ovipneumoniae* DNA copy number. Validation studies and fine mapping of markers from these genomic regions will allow for the possibility of applying selective breeding methods to reduce *M. ovipneumoniae* nasal detection from domestic sheep.

## Materials and Methods

### Ethics Statement

All animal care and handling procedures were reviewed and approved by the Washington State University Institutional Animal Care and Use Committee (Permit Number: 4885 and 4594) and/or by the U.S. Sheep Experiment Station Animal Care and Use Committee (Permit Numbers:15-04, 15-05).

### Sheep

Whole blood (jugular venipuncture) and nasal mucosal secretions were collected from ewes of Rambouillet (N = 321), Polypay (N = 222), and Suffolk (N = 104) breeds, ranging in age from 1 to 7 years. Repeated sampling occurred three times per year (February, April, and September/October) for 4 years (2015 – 2018). Ewes were managed as a single flock in an extensive rangeland production system. Sampling corresponded with mid-pregnancy (February), early-lactation (∼35 days post lambing, April), and post-weaning (∼30 days post-weaning, September/October) production stages. At mid-pregnancy and early-lactation sampling events, ewes had been managed in a penned, close-contact environment (e.g., dry-lot) and fed harvested feeds for at least 45 and 120 days, respectively. At the post-weaning sampling event, ewes had been managed in an extensive rangeland grazing environment. A total of 2,876 nasal swabs were collected for this study.

### *M. ovipneumoniae* Detection

Secretions were collected with sterile cotton swabs that were inserted approximately 10 cm into each nostril, consecutively, and rotated briefly. Swabs were stored without media at - 20°C until DNA was extracted with QiaAmp DNA Mini kit (Qiagen, Germantown, MD). Nasal shedding status of *M. ovipneumoniae* was determined by qPCR with species-specific primers 439F: TGATGGAACATTATTGCGCT and 439R: TGCCATTATTTGAAACRAGA. Reactions consisted of 1uL each (10uM) primer, 10uL of SsoFast EvaGreen Super mix (BioRad), 6uL of distilled water and 2uL of DNA template diluted 1:10. Each sample was run in triplicate on a CFX96 Touch system (BioRad, Hercules, CA) and quantified using a 10-fold dilution series standard curve ranging from 0 to 10^6^ copies. The standard curve was generated with DNA from the *M. ovipneumoniae* Y98 type-strain cultured in SP4 medium, pelleted by centrifugation and extracted with the QiaAmp DNA Mini kit.

### Genotyping

Blood was collected into EDTA-coated vacutainer tubes. Genomic DNA was isolated using the Invitrogen GeneCatcher gDNA 3–10 ml Blood Kit (Life Technologies, Carlsbad, CA) or Gentra PureGene (Qiagen, Germantown, MD), with minor modifications of manufacturers’ instructions. Quality and quantity of DNA were quantified using an ND-1000 spectrophotometer (Nanodrop, Wilmington, DE) and 300ng per ewe was sent for genotyping. Genotyping services were provided by Geneseek Inc. (Lincoln, NE) using the OvineHD BeadChip (Illumina Inc., San Diego, CA) with a set of greater than 600,000 SNP designed by the International Sheep Genome Consortium.

### Phenotype Analysis

Univariate analysis was conducted (SAS v9.4; SAS Inst. Inc., Cary, NC) to identify outliers and test for normality. *Mycoplasma ovipneumoniae* DNA copy number data were not normal, and therefore, values greater than zero were transformed to log_10_ to reduce heterogenous variances. To test if there were shedding differences a reduced mixed model in SAS that accounted for fixed effects of breed, age at sample collection, year of sample collection, and approximate month of sample collection, ewe as a repeated effect, and sire as a random effect was conducted. Tukey-Kramer adjustment was implemented to identify pair-wise differences.

### Association Analysis

A preliminary screen for high genotype call rates (>95%) was performed to select individuals for further analysis. Single nucleotide polymorphism inclusion screening criteria was conducted with PLINK1.9 as previously described [22]: missingness by individual (0.1), missingness by marker (0.05), minor allele frequency (0.01), and Hardy-Weinberg equilibrium (0.0000001, which corresponds to P = 0.05 after Bonferroni correction for 600,000 marker tests).

Quality control was performed on 647 individual samples and 606,006 SNP sites using Plink1.90 software. The following SNPs were excluded: 1) 203 for duplicate base pair location on the same chromosome; (2) 14,953 not in accordance with the Hardy–Weinberg equilibrium; 3) 31,770 with a minor allele frequency less than 1%; 4) 26,412 with genotyping rate less than 95%. In addition, multidimensional scaling was conducted to identify breed outliers. Three individuals were excluded because of failure to cluster with breed in multidimensional scaling (Figure 1). Genotyping rate was 99.8% for 644 sheep and 532,668 SNPs were used in the analyses.

**Figure 1.**
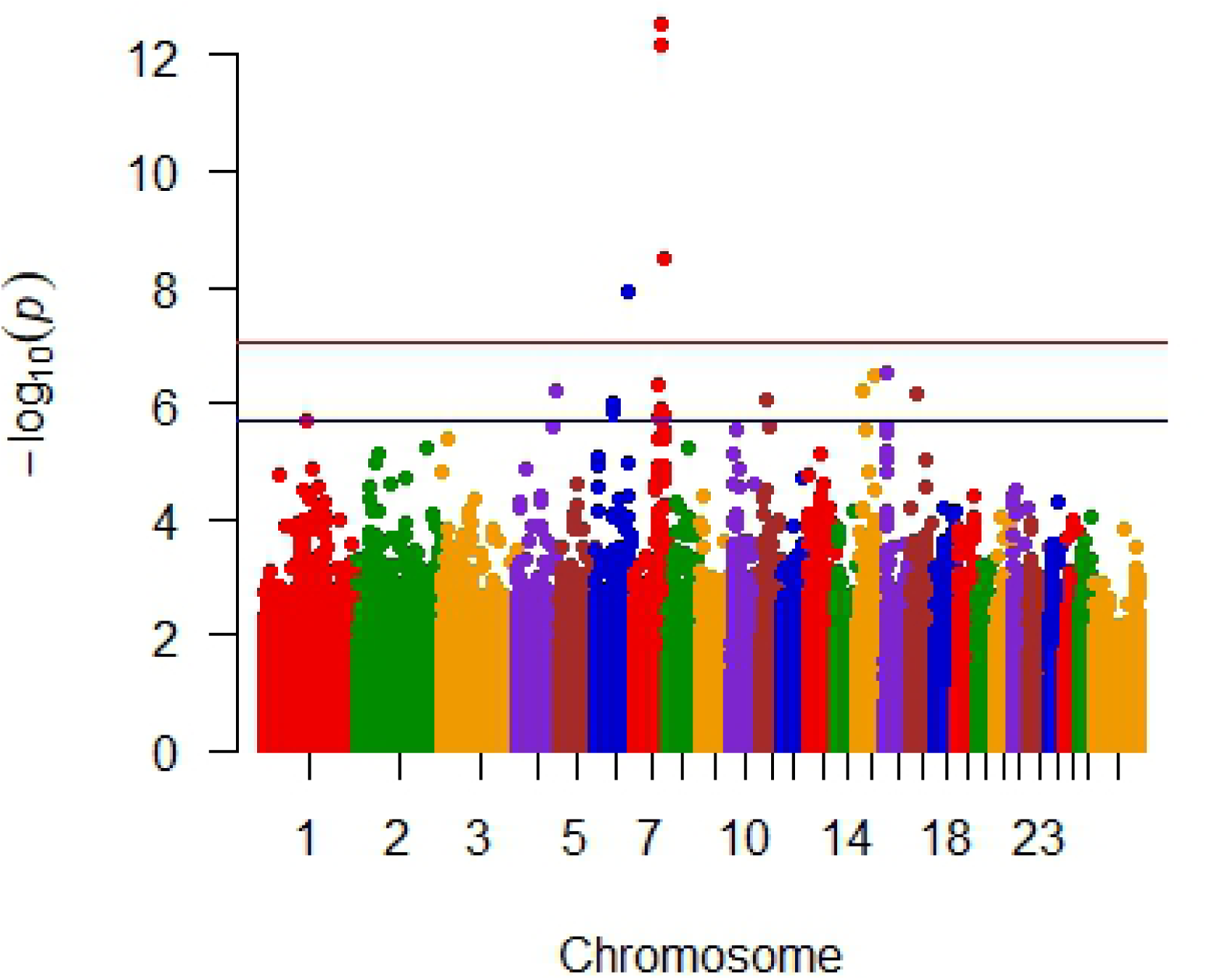
Plots of the first 2 principle components with 647 sheep showing breed outliers (A) and without the 3 outliers (B). Blue represents Rambouillet, pink represents Polypay, black represents Suffolk and green represents the 3 outliers.

Data were reformatted in PLINK1.9 for use with Efficient Mixed-Model Association eXpedited (EMMAX). The full model evaluated included fixed effects of breed or principle components 1 to 6 (eigenvalues greater than 4.9) and random effect of relationship matrix (Balding-Nichols estimate). For individual breed analysis, principle components with eigenvalues greater than 4.9 were Polypay PC1-2, Rambouilet PC1-4, and Suffolk PC1 were considered fixed effects and random effect of relationship matrix (Balding-Nichols estimate). Phenotypes considered were 1) binomial where all samples from an ewe were either greater or were less than or equal to100 DNA copy numbers, 2) categorical where a mixed detection among sampling times was added to the binomial data, and 3) untransformed mean DNA copy number. Genome-wide suggestive significance was set at a nominal P<9×10^−8^ consistent with other sheep GWAS using the OvineHD Beadchip [29]. In R, qqman [30] was used for visualization of results in Manhattan and quantile-quantile plots. Further, the significant SNPs were interrogated using a mixed model similar to those described above, accounting for log_10_ transformed *M. ovipneumoniae* DNA copy number, breed, age, year, sample time, and genotype with SAS.

Ensembl and NCBI were used to determine the location of the SNP within genome assembly OAR_Rambouillet_v1.0 as well as identify annotated genes within 100 kb of the SNP. The Illumina designated name of the SNP with the dbSNP rs# cluster id, the dataset where the significant SNP was identified, minor allele, and *M. ovipneumoniae* mean DNA copy trend associated with the minor allele can be found in Supplemental Table 1. Genotypes and phenotypes by ewe for the significant SNPs may be found at https://data.nal.usda.gov/dataset/data-genomic-regions-associated-mycoplasma-ovipneumoniae-presence-nasal-secretions-domestic-sheep. GeneCards and a search of Pubmed was used to provide known information about genes within 100kb of the associated SNPs. Reactome was used to determine the pathway(s) associated with the annotated genes.

**Table 1.** Mean *M. ovipneumoniae* log_10_ transformed DNA copy number in breed and age of ewes; year and month of sample collection.

|  |  | N | Mean | StdErr | P-value |
| --- | --- | --- | --- | --- | --- |
| <u>Breed</u> |  |  |  |  | <0.0001 |
|  | Polypay | 220 | 1.73<sup>a</sup> | 0.083 |  |
|  | Rambouillet | 320 | 1.12<sup>b</sup> | 0.071 |  |
|  | Suffolk | 104 | 1.97<sup>a</sup> | 0.12 |  |
| <u>Age (years)</u> |  |  |  |  | 0.0236 |
|  | 1 | 519 | 1.91<sup>a</sup> | 0.096 |  |
|  | 2 | 599 | 1.68<sup>b</sup> | 0.079 |  |
|  | 3 | 453 | 1.58<sup>b</sup> | 0.081 |  |
|  | 4 | 464 | 1.52<sup>b</sup> | 0.082 |  |
|  | 5 | 407 | 1.52<sup>b</sup> | 0.084 |  |
|  | 6 | 293 | 1.51<sup>b</sup> | 0.01 |  |
|  | 7 | 141 | 1.54<sup>a,b</sup> | 0.13 |  |
| <u>Year</u> |  |  |  |  | <0.0001 |
|  | 2015 | 370 | 1.22<sup>a</sup> | 0.086 |  |
|  | 2016 | 904 | 1.65<sup>b</sup> | 0.064 |  |
|  | 2017 | 974 | 1.6<sup>b</sup> | 0.066 |  |
|  | 2018 | 628 | 1.96<sup>c</sup> | 0.085 |  |
| <u>Month</u> |  |  |  |  | <0.0001 |
|  | February | 917 | 1.2<sup>a</sup> | 0.064 |  |
|  | April | 1137 | 1.91<sup>b</sup> | 0.062 |  |
|  | September/October | 822 | 1.71<sup>c</sup> | 0.068 |  |

a–c Means within Mean column for each effect with different superscripts differ ( $P < 0.05$ ). Mean separation tests were adjusted for multiple comparisons using the Tukey-Kramer option in PROC Mixed (SAS Inst. Inc., Cary, NC).

## Results

### *Mycoplasma ovipneumoniae* DNA Copy Number

Nasal mucosal secretions were collected from three to 10 times from each ewe, median number of collections was four. Untransformed *M. ovipneumoniae* DNA copy number ranged from 0 to 109,000 with a mean of 1,461 and median of 22. Transformed (log_10_) *M. ovipneumoniae* DNA copy number ranged from 0 to 5.03 with mean of 1.52 and median of 1.34. There were 224 ewes where DNA copy number was ≤ 100 (zero to low detection, considered not detected) ranging from three to eight sampling times with a mode of four. There were 94 ewes with greater than 100 DNA copy number (considered positive for each sample time) with a range of three to ten and a mode of four sampling times. Finally, there were 326 ewes with mixed levels of detection (above and below 100) when sampled over time with a range of three to six and a mode of four sampling times. Only 28 ewes had zero detect DNA copy number for all time points tested and ranged from three to six with a mode of four sampling times.

Breed of sheep significantly (P<0.0001) affected detected log_10_ transformed *M. ovipneumoniae* DNA copy number (Table 1). Rambouillet ewes had lower (P<0.0001) detected DNA copy numbers than Polypay and Suffolk ewes which were not different (P>0.21). Detected DNA copy number was greatest for yearlings and declined with increasing age until a slight increase at 7 years old (Table 1). Yearling ewes had significantly (P<0.046) higher detected DNA copy numbers than ewes 2 through 6 years of age. No other pair-wise age comparisons were different (P>0.08). Time during the year samples were collected significantly (P<0.0001) impacted DNA copy number (Table 1). All three sample periods were different (P<0.003) from one another where April had the highest and February had the lowest mean DNA copy number. Year of sample collection significantly (P<0.0001) impacted detected DNA copy number, with means increasing with time (Table 1). All years were different (P<0.004) in pair-wise comparisons except for 2016 and 2017 (P>0.87).

### Association Analysis

No significant genomic associations were identified when *M. ovipneumoniae* DNA copy number detection was considered binomial, where all samples from an ewe were either positive or undetected or when categorization included mixed detection. Seven SNPs were identified as significantly (P<10^−8^) associated with untransformed mean *M. ovipneumoniae* nasal shedding (Figure 2). These SNPs represented 3 regions on chromosomes 6 and 7 (Table 2) where the least frequent genotype for all seven SNPs had the highest log_10_ DNA copy number and the minor allele was present in all breeds. There were 4 closely spaced SNPs on chromosome 7, approximately 21Kb apart, that were highly significant (2.72 to 6.60 × 10^−13^) within 50Kb of *Transmembrane protein 229B* (*TMEM229B*). An additional SNP, at approximately 91Kb, on chromosome 7 was significant at 3.15 × 10^−9^ and was within 50Kb of a postulated gene *ENSOARG00000026723*. The two SNPs on chromosome 6 were in introns of *Protein Tyrosine Phosphatase Non-Receptor Type 13* (*PTPN13*; 1.21 × 10^−8^). Quantile-Quantile plots are shown in Figure 3, where A includes all SNPs in the dataset analyzed, showing extreme deviation with the most significant SNPs. Panel B in Fig. 3 shows a Q-Q plot that had the top seven SNPs included in the model when reanalyzed, demonstrating their impact.

**Table 2.** Significant SNPs associated with untransformed *M. ovipneumoniae* mean DNA copy number in the all-breed analysis. Listed chromosome position is from Oar_Rambouillet v1.0.

| SNP | CHR | Position | MAF (%) | Nominal P-value | Closest gene |
| --- | --- | --- | --- | --- | --- |
| rs159990985 | 6 | 111960643 | 3.261 | 1.21E-08 | PTPN13 |
| rs159991022 | 6 | 111973304 | 3.261 | 1.21E-08 | PTPN13 |
| rs426816169 | 7 | 82832532 | 3.571 | 2.72E-13 | TMEM229B |
| rs411153659 | 7 | 82840000 | 3.571 | 2.72E-13 | TMEM229B |
| rs400632990 | 7 | 82852837 | 3.339 | 6.60E-13 | TMEM229B |
| rs427671730 | 7 | 82854206 | 3.339 | 6.60E-13 | TMEM229B |
| rs424565925 | 7 | 99167837 | 4.51 | 3.15E-09 | ENSOARG00000026723 |

**Figure 2.**
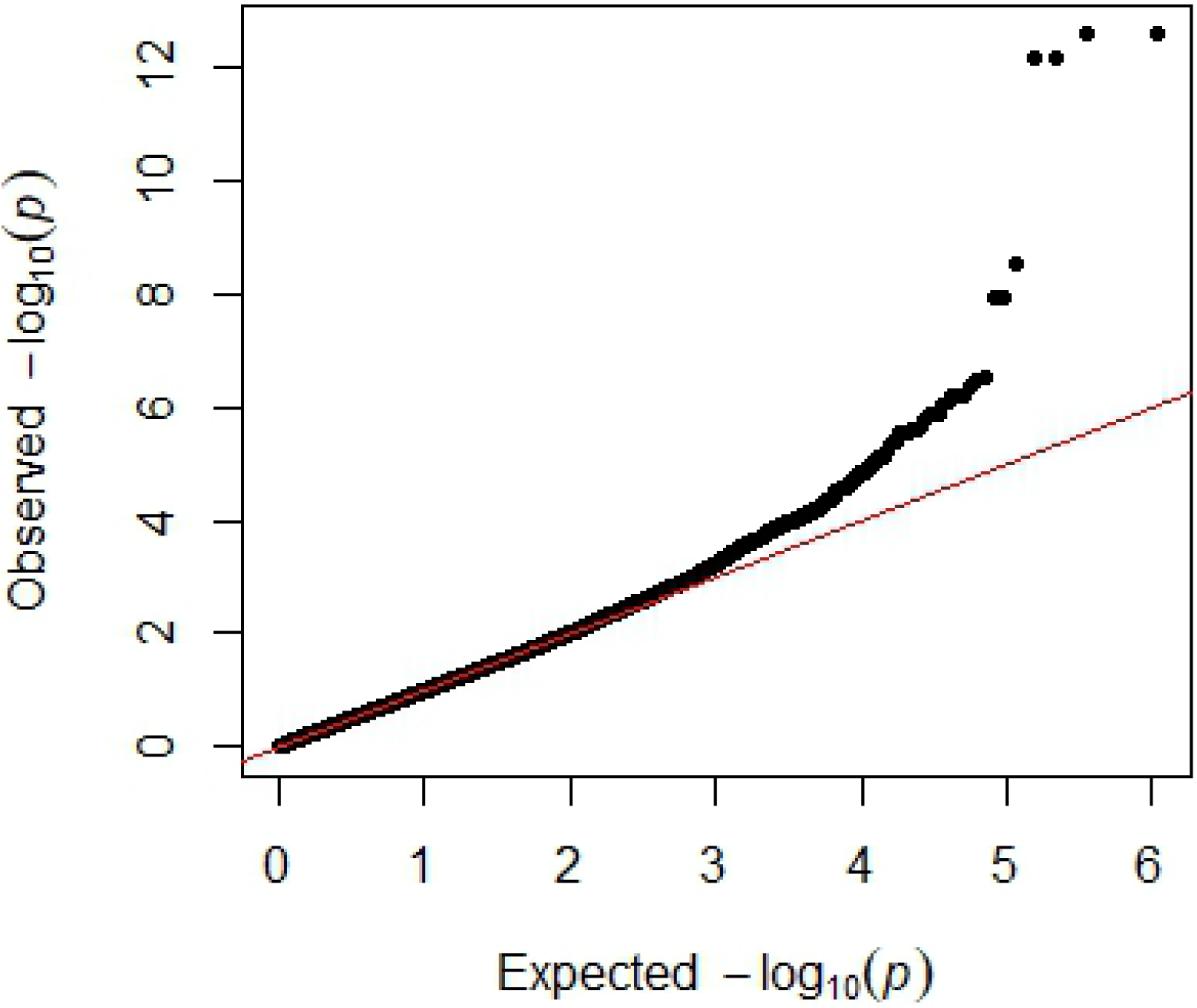
Manhattan plot of all breed analysis with untransformed *M. ovipneumoniae* mean DNA copy number showing significant SNPs on chromosomes 6 and 7. Genome significant line is red and genome suggestive line is blue.

**Figure 3.**
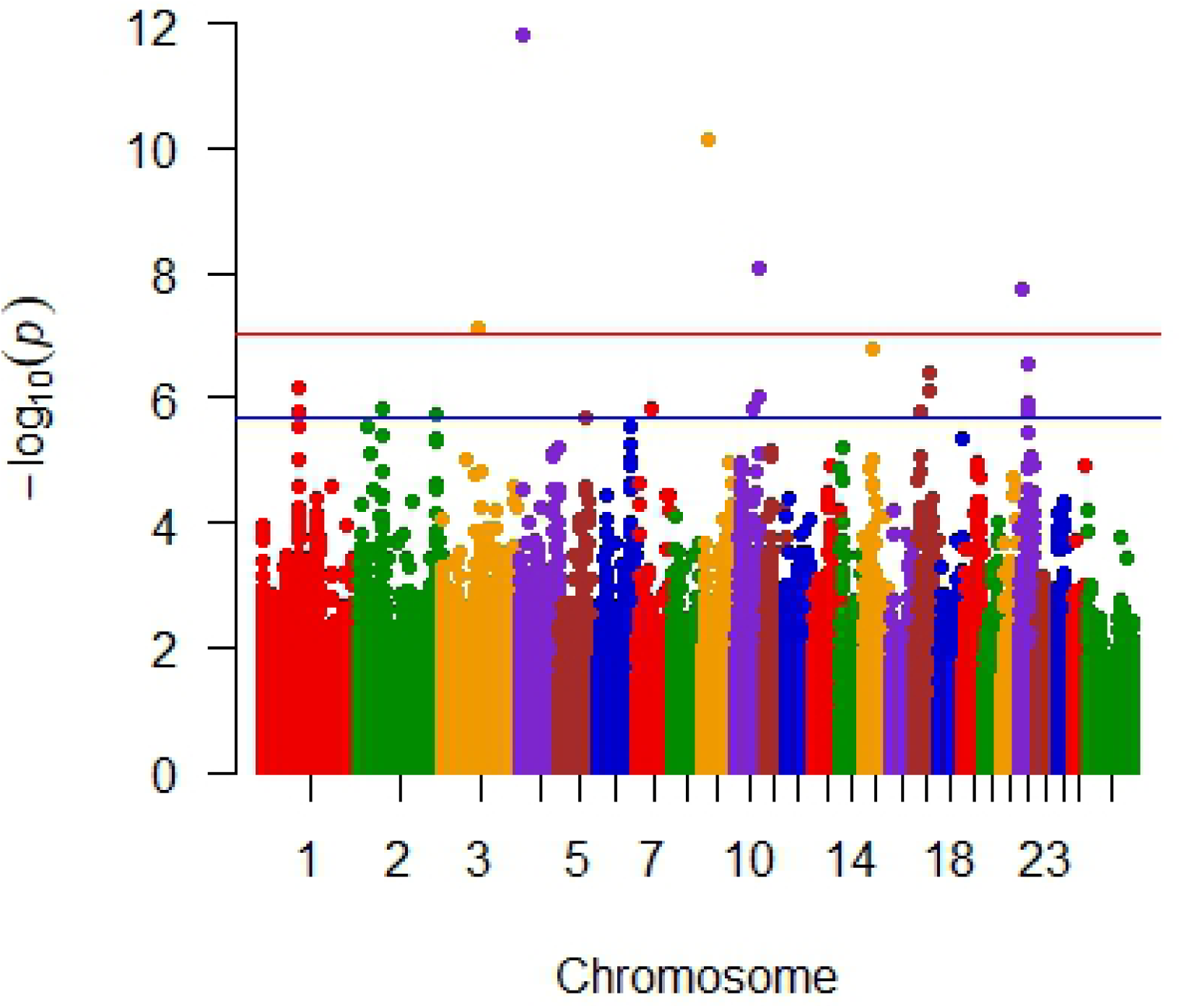
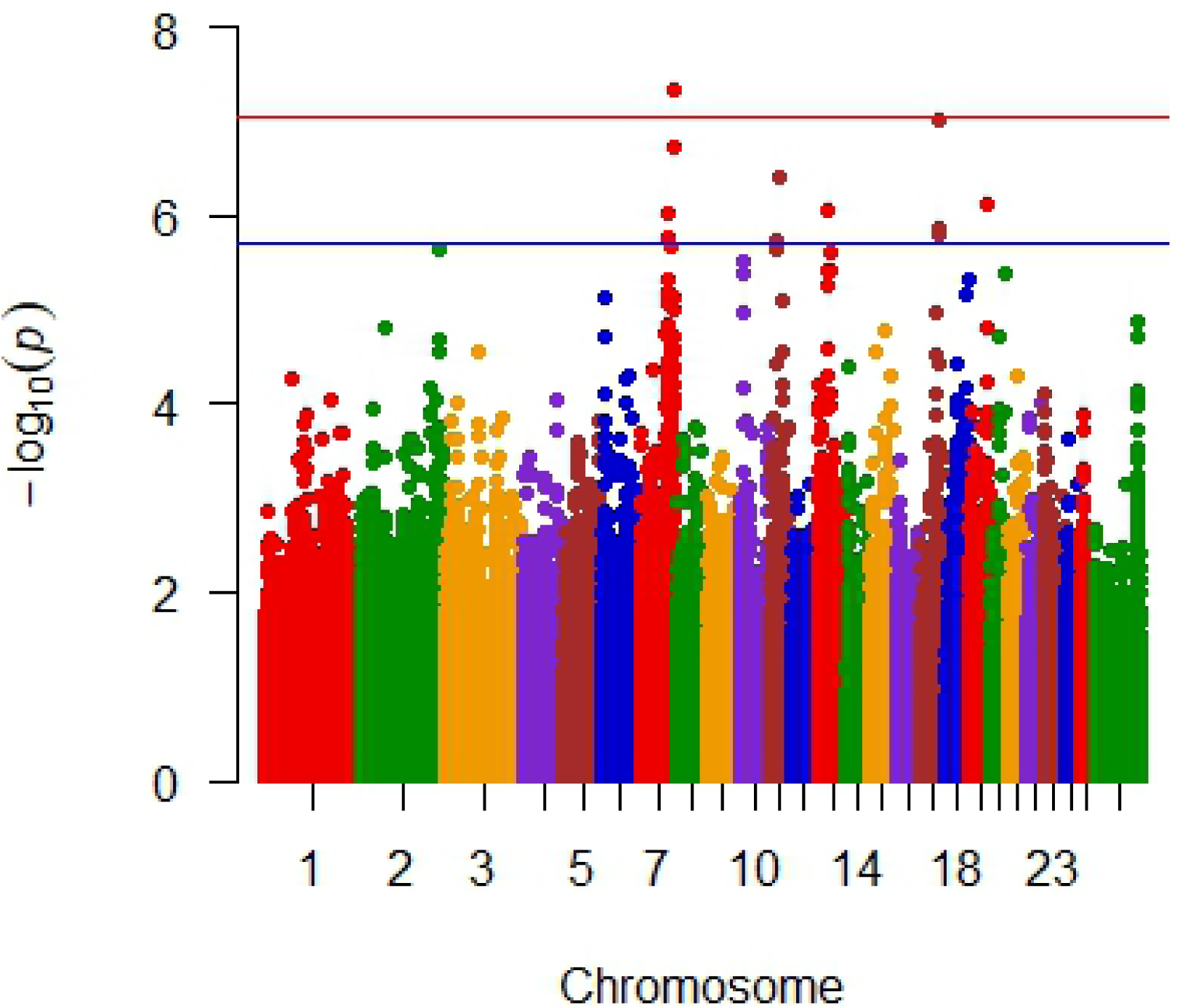
Quantile-Quantile plots for the all-breed analysis with untransformed *M. ovipneumoniae* mean DNA copy number, showing deviation due to significant SNPs (A) and less deviation when top 7 SNPs are included as covariates in the analysis (B).

Analysis of untransformed *M. ovipneumoniae* DNA copy number was conducted within each breed with 320, 220, and 104 ewes classified as Rambouillet, Polypay, and Suffolk, respectively (Table 3). Within Rambouillet, there were nine SNPs on five chromosomes (3, 4, 9, 10, and 22) that were significantly (<10^−8^) associated with *M. ovipneumoniae* mean DNA copy number (Figure 4). Three significant SNPs on chromosome 4 were within *Thyrotropin Releasing Hormone Degrading Enzyme* (*TRHDE*) and one significant SNP was within 25Kb of *TRHDE*. The two significant SNPs on chromosome 22 were within *Solute Carrier Family 16 Member 12* (*SLC16A12*). Analysis of only Polypay identified three SNPs on two chromosomes (7 and 17) that were significantly (P<4.63×10^−8^ and 9.34×10^−8^, respectively) associated with *M. ovipneumoniae* mean DNA copy number (Figure 5). The significant SNPs on chromosome 17 were within *Lysine-specific demethylase 2b* (*KDM2B*) and *Anaphase Promoting Complex Subunit 7* (*ANAPC7*). Nine SNPs on two chromosomes (7 and 15) were significantly associated with *M. ovipneumoniae* mean detection in the Suffolk-only analysis (Figure 6). Eight of these nine significant SNPs were on chromosome 15, with three of the SNPs within 31Kb of one another and two SNPs within *piwi (P element-induced Wimpy)-like RNA-mediated gene silencing 4* (*PIWIL4*). In addition, one SNP was within *Unc-13 Homolog C* (*UNC13C*) on chromosome 15 and one on chromosome 7 was within 10Kb of *Cullin 5* (*CUL5*).

**Table 3.** Significant SNPs associated with untransformed *M. ovipneumoniae* mean DNA copy number in the individual breed analysis. Listed chromosome position is from Oar_Rambouillet v1.0.

| SNP | CHR | Position | MAF(%) | Breed | Nominal P-value | Closest Gene |
| --- | --- | --- | --- | --- | --- | --- |
| rs403152750 | 3 | 115055558 | 1.56 | Rambouillet | 5.30E-08 | TRHDE |
| rs404662185 | 3 | 115060178 | 1.57 | Rambouillet | 5.24E-08 | TRHDE |
| rs409357561 | 3 | 115062407 | 1.56 | Rambouillet | 5.30E-08 | TRHDE |
| rs407012017 | 3 | 115135930 | 1.56 | Rambouillet | 5.30E-08 | TRHDE |
| rs398278706 | 4 | 10400045 | 1.25 | Rambouillet | 8.31E-13 | ENSOARG00000025219 |
| rs420722286 | 7 | 57732050 | 5.34 | Suffolk | 7.23E-09 | UNC13C |
| rs402243563 | 7 | 99714676 | 1.406 | Polypay | 4.63E-08 | N/A |
| rs160633679 | 9 | 15144452 | 1.122 | Rambouillet | 1.43E-10 | ENSOARG00000016023 |
| rs429475222 | 10 | 75915173 | 1.406 | Rambouillet | 6.81E-09 | N/A |
| rs410915115 | 15 | 17401773 | 1.923 | Suffolk | 3.53E-10 | PIWIL4 |
| rs406526355 | 15 | 17412929 | 1.923 | Suffolk | 3.53E-10 | PIWIL4 |
| rs413355105 | 15 | 17431988 | 3.365 | Suffolk | 4.79E-08 | PIWIL4 |
| rs406613692 | 15 | 17480590 | 2.885 | Suffolk | 4.82E-09 | ENSOAG0000009780 |
| rs418601541 | 15 | 19401648 | 2.885 | Suffolk | 2.02E-08 | CUL5 |
| rs590224797 | 15 | 21333112 | 4.327 | Suffolk | 2.92E-08 | C11of87 |
| rs406524055 | 15 | 21333678 | 4.327 | Suffolk | 2.92E-08 | C11of87 |
| rs409884491 | 15 | 21442130 | 2.404 | Suffolk | 1.27E-10 | C11of87 |
| rs402253048 | 17 | 62654668 | 1.591 | Polypay | 9.34E-08 | KDM2B |
| rs160856157 | 17 | 63273726 | 1.591 | Polypay | 9.34E-08 | ANAPC7 |
| rs429858348 | 22 | 12903174 | 1.406 | Rambouillet | 1.31E-08 | SLC16A12 |
| rs413436283 | 22 | 12912440 | 1.406 | Rambouillet | 1.31E-08 | SLC16A12 |

**Figure 4.**
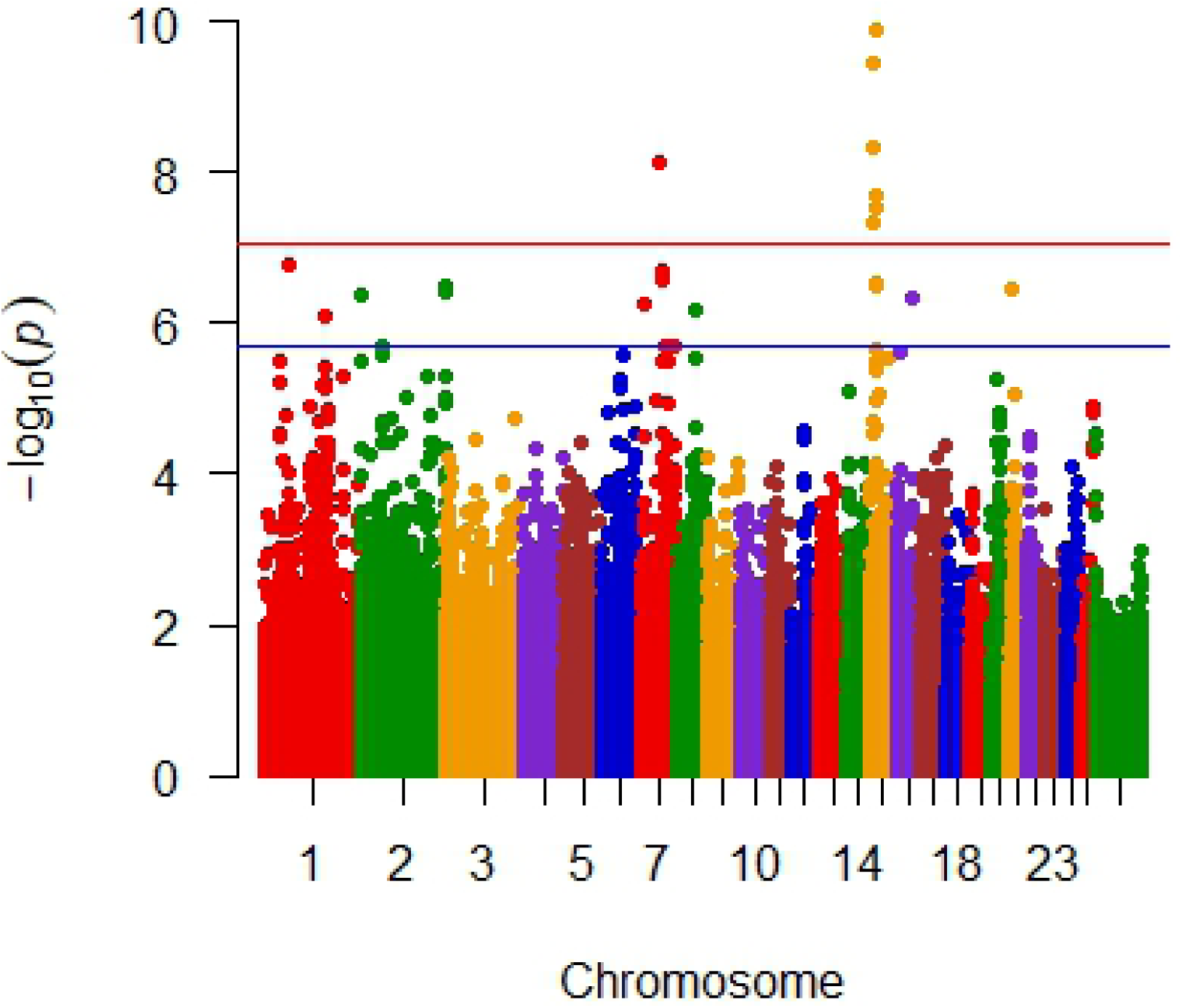
Manhattan plot of Rambouillet analysis with untransformed *M. ovipneumoniae* mean DNA copy number. Significant SNPs are on chromosomes 3, 4, 9, 10, and 22. Genome significant line is red and genome suggestive line is blue.

**Figure 5.**
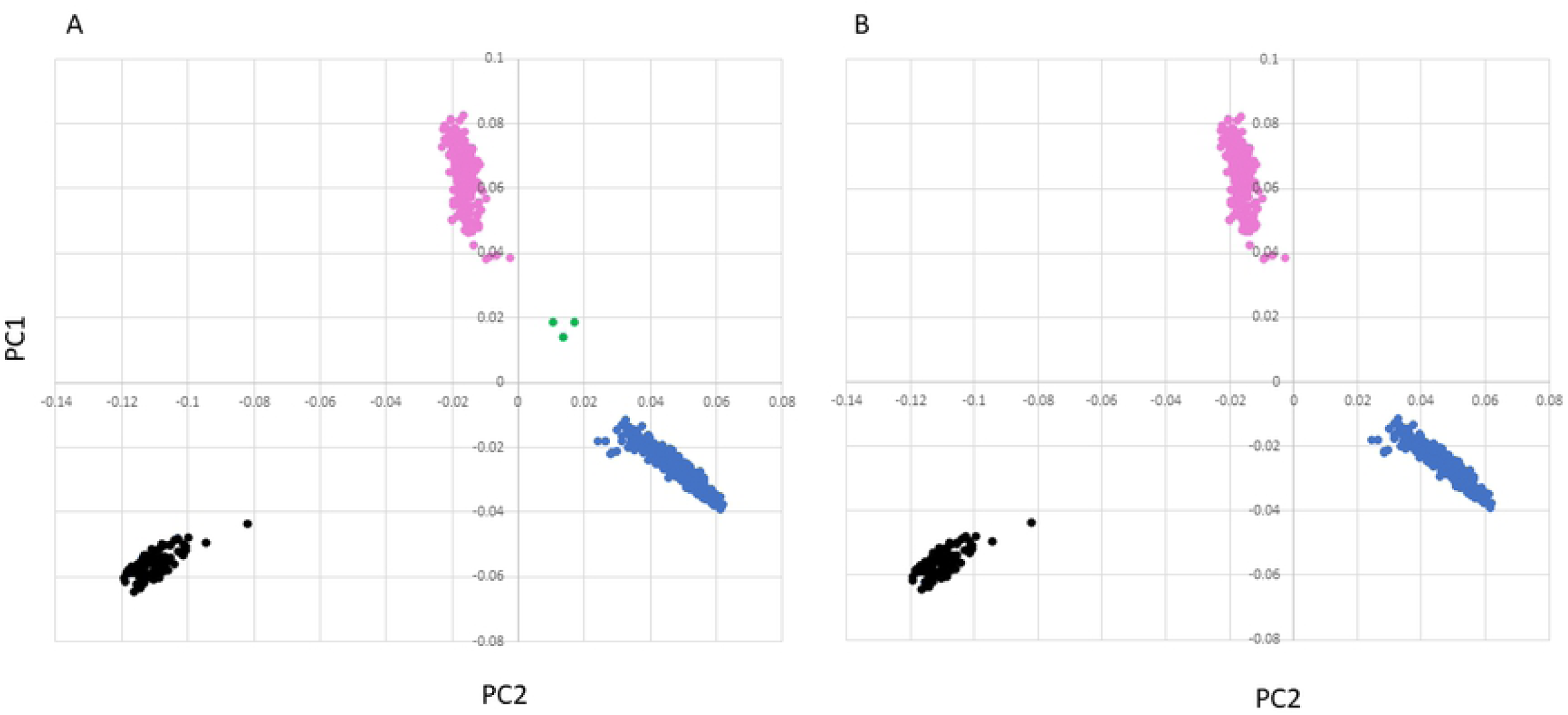
Manhattan plot of Polypay analysis with untransformed *M. ovipneumoniae* mean DNA copy number. Significant SNPs are on chromosomes 7 and 17. Genome significant line is red and genome suggestive line is blue.

**Figure 6.**
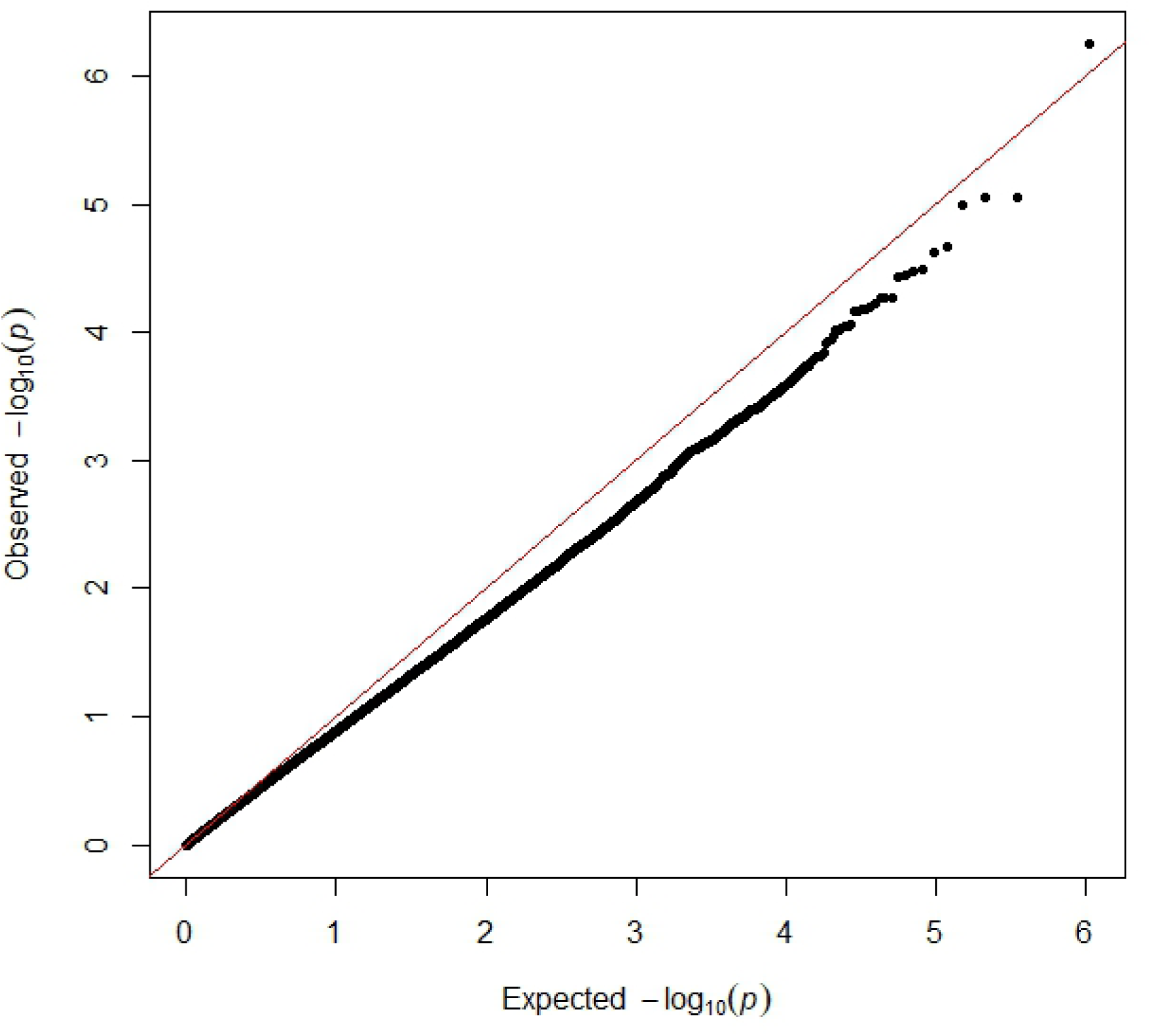
Manhattan plot of Suffolk analysis with untransformed *M. ovipneumoniae* mean DNA copy number. Significant SNPs are on chromosomes 7 and 15 Genome significant line is red and genome suggestive line is blue.

## Discussion

This is the first known publication of repeated, sequential *M. ovipneumoniae* nasal secretion testing in domestic sheep, with testing spanning 4 contiguous years. Detection and quantification of *M. ovipneumoniae* was influenced by sheep breed and age, and sampling time and year of sample collection. In concordance with Cheng et al. [11], we identified differences in the detection of *M. ovipneumoniae* in three different breeds of sheep. Rambouillet had the lowest detected DNA copy numbers for all sampling times measured. Similar studies that found Rambouillet had lower levels of ovine progressive pneumonia virus compared with Polypay and Columbia [22, 31]. While not significantly different from Polypay, Suffolk had numerically the highest mean detection of *M. ovipneumoniae*. A Japanese survey study found variable detection (0 to 53%) in 13 different flocks from three prefectures and the positive flocks had at least one serologically positive animal representing 9 different breeds and crossbreds, including Suffolk [32]. Flocks with greater than 150 sheep or Suffolk tended to have a higher percentage of positive sheep [32] which this study agrees with and supports.

We identified a shedding pattern among the sheep, where a subset consistently did not have detectible *M. ovipneumoni*ae DNA, was always detected, or had mixed detection. Understanding this variability is important for conducting genomic studies and improves the possibility for selection of non-shedding sheep. Unfortunately, there were too few animals in the always detected category to identify genomic associations with the categorical analysis in this study. In other livestock, approximately 4.8% of cattle were positive (nasal secretions) for *Mycoplasma bovis* testing [33]. A longitudinal study of dairy cattle also found a low positivity rate of about 4% but identified one heifer that was shedding *M. bovis* for each time tested over 2 years [34]. In this study, we found 14.6% of ewes were positive every time tested and an overall positivity rate of 38.3%. Detection of *M. ovipneumoniae* was greatest in yearlings and decreased as the sheep aged. Converse of DNA copy number data, serological data indicated the number of positive animals increased with age [32] which might suggest that some individuals seroconvert at extended intervals post-exposure.

April had the greatest mean DNA copy number. When sheep are in close confinement, such as for the longest period before testing in April, opportunity for direct-contact transmission increases. Experimental transmission of *M. hyopneumoniae* in pigs has been demonstrated to be due to direct contact and less effective with indirect exposure [35]. September/October mean DNA copy number was intermediate to February and April. This was somewhat unexpected as the sheep were on open range grazing for approximately 3 months before testing. Although, while on open range, sheep are herded into a small area with close contact every night to better protect them from predators. Temperature and humidity were found to impact *Mycoplasma* infectivity in a Spanish study of lambs, with higher temperature and lower humidity having greater rates of infection [36]. Specifically, they found February, March, and September to October had the lowest detection and April through August had the greatest. The Spanish and this study together demonstrate that transmission of *M. ovipneumoniae* among sheep is not necessarily just a function of proximity, but that climate may also be a factor.

Genomic associations with *M. ovipneumoniae* detection were identified in all-breed and individual-breed analyses. Within the all-breed analysis, significant SNPs were close to *TMEM229B* on chromosome 7 and *PTPN13* on chromosome 6. Both these genes are found in immune pathways, and impact T cells and innate lymphoid cell type 2. Specifically, *TMEM229B* was found to be down regulated in memory CCR6+ T cells in an HIV study [37]. The CCR6+ T cells direct Th17 cells to effector mucosal sites [38, 39], where they secrete cytokines. Some of these cytokines stimulate neutrophils and other immune cells to promote clearance of extracellular bacteria, such as *Streptococcus pyogenes* (*Sp*), *Klebsiella pneumoniae*, and *Bordetella pertussis* [40–45]. Interestingly, *TMEM229B* was identified as under extreme selection in chickens from tropical and arid climates [46]. The immune system of these chickens would be hypothesized to respond differently to pathogens as the two environments in the signatures of selection study were vastly different. While *PTPN13* has been shown to play a role in T cell activation [47], it is specifically expressed in Th17 T cells [48] and has variable expression in T cells of people with Rett Syndrome [49]. In addition, *PTPN13* is a conserved transcript in innate lymphoid cell type 2 (ILC2), which are considered helper-like and found in upper and lower airway and blood of humans [50] and mice [51]. In mouse lung, ILC2s were found to proliferate upon intranasal application of allergens, and memory-like lung ILC2s displayed upregulated expression of cytokines and their respective receptors, but also receptors for alarmins (*IL25R, ST2*), and genes associated with cell activation and proliferation (*BCL2A1B, IER3*) [51]. Also, *PTPN13* (PTP-BL) interacts with the Rho effector *kinase protein kinase C-related kinase 2* (*PRK2*) [53] and *PRK2* has been demonstrated to have a role in cilia signaling [54] as well as regulation of the formation of apical junctions in bronchial epithelial cells [55]. Infection of a murine macrophage cell line with *M. ovipneumoniae* increases levels of *nucleotide-binding oligomerization domain-containing 2* (*NOD2*) and *NOD2* along with activation of c*-Jun NH2-terminal protein kinase* (*JNK*) induces autophagy [56]. *NOD2* interacts with *RAC family small GPTase 1* (*Rac1*) in membrane ruffles [57] and it has been suggested that Rho GTPases such as *Rac1* regulate *PRK2* activity during control of the cell cycle [58].

In individual-breed analyses, 21 SNPs on 8 chromosomes were identified as associated with mean *M. ovipneumoniae* detection. One of these SNPs was in a similar chromosomal region as the all-breed analysis, but the majority were on different chromosomes. Specifically, eight SNPs were identified on chromosome 15 within three regions in Suffolk. *PIWIL4*, on chromosome 15, has been identified in the chromatin pathway [59, 60], where human *PIWIL4* has a role in regulating gene expression in immune cells with gene silencing [61]. *In vitro* experiments showed that PIWIL4 knockdown in primary human cells from people with HIV-1 on suppressive combined antiretroviral therapy had increased HIV-1 transcription and reversed HIV-1 latency in HIV-1 latently infected Jurkat T cells and primary CD4+ T lymphocytes, as well as resting CD4+ T lymphocytes [62]. Mutations in *CUL5*, also located on chromosome 15, have been associated with more rapid loss of CD4+ T cell in HIV infected patients [63].

Two SNPs on chromosome 17, near two different genes, were significantly associated in the Polypay analysis. The expression of *KDM2B* has been shown to help maintain ILC3 cells, while reduction in ILC3s leads to susceptibility to bacterial infection [64]. Interestingly, optimal Th17 cell formation in mice after *Streptococcus pyogenes* infection requires polycomb repressive complex 1.1, which includes *KDM2B* [65]. The other significant gene, *ANAPC7*, is a part of the anaphase promoting complex/cyclosome (APC/C), which is a substantial E3 ubiquitin ligase that controls mitosis by catalyzing ubiquitination of key cell cycle regulatory proteins [66]. In human lungs, an overexpression of *ANAPC7* protein has been identified and differential expression was found between smokers and non-smokers [67].

In the Rambouillet analysis, four SNPs were within or very near *TRHDE* on chromosome 3. This is an important regulator of thyroid hormones [68]. Although there does not appear to be a direct involvement of this gene with the immune system, a study found T- and B-cells from hyperthyroidic mice had higher mitogen-induced proliferation compared with the same cells from hypothyroidic mice [69]. Two SNPs were near an annotated gene, *SLC16A12*, a transmembrane transporter on chromosome 22. This gene was found to be induced by budesonide (a corticosteroid to treat asthma by reducing inflammation) in A549, BEAS-2B and primary human bronchial epithelial cells [70]. Interestingly, another GWAS examining susceptibility of mice to *Streptococcal pneumoniae* infection found an association with *Slc16a12* as well and the authors made the correlation with *Slc11a1*, another known solute carrier transporter, that has been found to affect susceptibility to bacterial infections [71].

## Conclusion

Domestic sheep nasal mucosal secretions had variable mean *M. ovipneumoniae* DNA copy number as detected with a new species-specific qPCR. Rambouillet had the lowest mean while Suffolk and yearlings had the greatest. Time of year samples were collected affected detected DNA copy number. Twenty-eight SNPs on 9 different chromosomes were associated with mean DNA copy number. Annotated genes near these SNPs have been identified in immune function pathways (*TMEM229B, PTPN13, CUL5*, and *ANAPC7*), chromatin organization (*KDM2B*), and gene silencing (*PIWIL4*). Nasal shedding of *M. ovipneumoniae* may affect mucosal immunity, specifically Th17 cells. This data will be used to identify causal mutations that then may be developed into commercial markers for the domestic sheep industry.

## Acknowledgements

The authors thank Nic Durfee for sample handling and generation of qPCR data, Caylee Birge and Codie Durfee for sample handling and Ralph Horn, James Allison, Lori Fuller, and ADRU staff for technical assistance. The authors thank Dr. Maggie Highland for participating in concept development, sample coordination and collection, qPCR results analysis, and technical consultation. We also acknowledge current and former USDA, Range Sheep Production Efficiency Research Unit (Dubois, ID, USA) employees Tom Kellom and Natalie Pierce for data collection, curation, and management. We gratefully appreciate Shiquan Wang, Mark Williams, Boyd Leonard, Ella Ybarlucea, Harley Carpenter, Joel Billman, Brad Eddins, Lyn Mortensen, Jack Hensley, Nicole Strong, and Jennifer Barnett for animal management, husbandry, and sample-collection assistance.

---

**Supplemental Table 1.** Name of significant SNPs on OvineHD Beadchip, rs number, the animal set in which the SNPs were identified, the minor allele, and trend of *M. ovipneumoniae* transformed log_10_ mean DNA copy number associated with minor allele.

| Illumina SNP name | dbSNP rs# cluster id | Animal Set | Minor Allele | Trend |
| --- | --- | --- | --- | --- |
| oar3_OAR6_101266504 | rs159990985 | All | C | Higher |
| oar3_OAR6_101279171 | rs159991022 | All | T | Higher |
| oar3_OAR9_13890848 | rs160633679 | Rambouillet | A | Higher |
| oar3_OAR17_53882329 | rs160856157 | Polypay | C | Higher |
| oar3_OAR4_8831863 | rs398278706 | Rambouillet | A | Higher |
| oar3_OAR7_76225469 | rs400632990 | All | T | Higher |
| oar3_OAR7_91908439 | rs402243563 | Polypay | A | Higher |
| oar3_OAR17_53261568 | rs402253048 | Polypay | G | Higher |
| oar3_OAR3_108633720 | rs403152750 | Rambouillet | A | Higher |
| oar3_OAR3_108638337 | rs404662185 | Rambouillet | G | Higher |
| oar3_OAR15_19011353 | rs406524055 | Suffolk | T | Higher |
| OAR15_15503325.1 | rs406526355 | Suffolk | C | Higher |
| oar3_OAR15_15453723 | rs406613692 | Suffolk | T | Higher |
| oar3_OAR3_108714118 | rs407012017 | Rambouillet | C | Higher |
| oar3_OAR3_108640566 | rs409357561 | Rambouillet | G | Higher |
| oar3_OAR15_18879871 | rs409884491 | Suffolk | A | Higher |
| oar3_OAR15_15371024 | rs410915115 | Suffolk | T | Higher |
| oar3_OAR7_76212720 | rs411153659 | All | A | Higher |
| oar3_OAR15_15401315 | rs413355105 | Suffolk | T | Higher |
| oar3_OAR22_10938408 | rs413436283 | Rambouillet | A | Higher |
| oar3_OAR15_17220107 | rs418601541 | Suffolk | G | Higher |
| oar3_OAR7_53135906 | rs420722286 | Suffolk | G | Higher |
| oar3_OAR7_91281519 | rs424565925 | All | T | Higher |
| oar3_OAR7_76205432 | rs426816169 | All | T | Higher |
| oar3_OAR7_76226838 | rs427671730 | All | G | Higher |
| oar3_OAR10_67992867 | rs429475222 | Rambouillet | G | Higher |
| oar3_OAR22_10929142 | rs429858348 | Rambouillet | T | Higher |
| oar3_OAR15_19011908 | rs590224797 | Suffolk | T | Higher |

